# Proteome profiling of cerebrospinal fluid using machine learning shows a unique protein signature associated with APOE4 genotype

**DOI:** 10.1101/2024.04.18.590160

**Authors:** Artur Shvetcov, Shannon Thomson, Ann-Na Cho, Heather M. Wilkins, Joanne H. Reed, Russell H. Swerdlow, David A. Brown, the Alzheimer’s Disease Neuroimaging Initiative, Caitlin A. Finney

**Affiliations:** Department of Psychological Medicine, Sydney Children’s Hospital Network, Sydney, NSW, Australia; Neuroinflammation Research Group, Centre for Immunology and Allergy Research, Westmead Institute for Medical Research, Sydney, NSW, Australia; School of Medical Sciences, Faculty of Medicine and Health, The University of Sydney, Sydney, NSW, Australia; Human Brain Microphysiology Systems Group, School of Biomedical Engineering, Faculty of Engineering, The University of Sydney, Sydney, NSW, Australia; University of Kansas Alzheimer’s Disease Centre, Kansas City, KS, USA; Department of Biochemistry and Molecular Biology, University of Kansas Medical Centre, Kansas City, KS, USA; Department of Neurology, University of Kansas Medical Centre, Kansas City, KS, USA; Autoimmunity and Amyloidosis Research Group, Centre for Immunology and Allergy Research, Westmead Institute for Medical Research, Sydney, NSW, Australia; Department of Molecular and Integrative Physiology, University of Kansas Medical Centre, Kansas City, KS, USA; Department of Immunopathology, Institute for Clinical Pathology and Medical Research-New South Wales Health Pathology, Sydney, NSW, Australia; Westmead Clinical School, Faculty of Medicine and Health, The University of Sydney, Sydney, NSW, Australia

**Keywords:** APOE4, Alzheimer’s disease, cerebrospinal fluid, proteomics, machine learning

## Abstract

**INTRODUCTION:** Proteome changes associated with APOE4 variant carriage that are independent of Alzheimer’s disease (AD) pathology and diagnosis are unknown. This study investigated APOE4 proteome changes in people with AD, mild cognitive impairment, and no impairment.

**METHODS:** Clinical, APOE genotype, and cerebrospinal fluid (CSF) proteome and AD biomarker data was sourced from the Alzheimer’s Disease Neuroimaging Initiative (ADNI) database. Proteome profiling was done using supervised machine learning.

**RESULTS:** We found an APOE4-specific proteome signature that was independent of cognitive diagnosis and AD pathological biomarkers, and increased risk of progression to cognitive impairment. Proteins were enriched in brain regions including the caudate and cortex and cells including endothelial cells, oligodendrocytes, and astrocytes. Enriched peripheral immune cells included T cells, macrophages, and B cells.

**DISCUSSION:** APOE4 carriers have a unique CSF proteome signature associated with a strong brain and peripheral immune and inflammatory phenotype that likely underlies APOE4 carriers’ vulnerability to cognitive decline and AD.

## 1. BACKGROUND

A mutation in the apolipoprotein E gene called ε4 (APOE4) is the single biggest genetic risk factor for late onset Alzheimer’s disease (AD) accounting for between 40-60% of AD’s genetic variability ^1,2^. ApoE itself is involved in lipid transport in the plasma and central nervous system and the ApoE4 isoform has been linked to changes in stability that may contribute to protein misfolding (and the misfolded protein response), aggregation, and proteolytic fragmentation ^3^. APOE4 carriers with AD typically have an onset between 2-10 years earlier than their non-carrier counterparts depending on the number of alleles ^2^. Despite the clear relationship between APOE4 and AD, we still have a poor understanding of the underlying mechanisms.

To date, most research has relied on the use of animal models to study the pathomechanisms underlying APOE4 carriage that may have limited translatability to human AD. For example, murine ApoE does not show N- and C-terminal domain interactions ^3,4^ and there are APOE gene promoter differences between mice and humans ^5^. Even though murine ApoE can be modified to reproduce some known metabolic properties of human apoE ^3,4^, or be replaced entirely by human ApoE ^6^, it’s not clear how these changes affect the biological system and downstream molecular interactions within it. Further, the use of familial AD mice that express known causal genetic variants in amyloid precursor protein (APP), presenilin 1 (PSEN1) or PSEN2 models only a minority of early onset, familial AD, with unclear implications for modeling the more common late onset AD ^7^.

Relative to animal model studies, there are fewer human studies examining potential mechanisms underlying APOE4 carriers’ increased AD risk. Previous research has shown that cognitively unimpaired adult APOE4 carriers have changes to cognitive function and memory ^8^ accompanied by structural and functional brain differences relative to non-carriers. This includes, for example, decreases in hippocampal volume ^9^, lower neurite density in the entorhinal cortex ^10^, changes in brain co-activation networks ^11–13^, and evidence of higher brain amyloid deposition and tau ^14–16^. Human stem cell-derived astrocytes from APOE4 homozygous carriers were found to be less able to promote neuronal survival and synaptogenesis ^17^. Stem cell-derived APOE4 neurons have found enhanced synthesis and intracellular signaling, including via MAP kinase ^18,19^, degeneration of GABAergic neurons ^20^, higher levels of tau phosphorylation ^20^, and increased Aβ production ^19,20^. Similarly, stem cell-derived pericytes and blood brain barrier (BBB) models show APOE4 leads to increased amyloid accumulation and dysregulated nuclear factor of activated T cells (NFAT) signaling^21^. Further demonstrating the importance of BBB dysfunction, a recent study found that APOE4 carriers, even those who are cognitively unimpaired, have an increased breakdown of the BBB in the hippocampus and medial temporal lobe linked to pericyte injury and activation of the BBB-degrading cyclophilin A-matrix metalloproteinase-9 pathway ^22^. In line with a BBB phenotype, recent evidence suggests that APOE4 carriers with AD have differences in cerebrospinal fluid (CSF) and plasma proteins relative to non-carriers with AD ^23–26^. These studies, however, have been limited by including only a small number of proteins (<300) and examining protein changes through the lens of brain-specific AD pathology including aggregated amyloid-β. Proteome-wide changes independent of AD brain pathology and whether these extend to APOE4 carriers irrespective of cognitive status remains unknown, limiting our understanding of whether these CSF proteome changes underly APOE 4 carriers’ vulnerability to AD.

To address this, we use a combination of machine learning and functional enrichment analyses to profile the CSF proteome of APOE4 carriers with and without cognitive impairment from the Alzheimer’s Disease Neuroimaging Initiative (ADNI) cohort.

## 2. METHODS

### 2.1. Data and Participants

We used clinical, APOE genotype, and CSF proteome data generated from the Alzheimer’s Disease Neuroimaging Initiative (ADNI) cohort for this study. All data is accessible through the ADNI database at (https://ida.loni.usc.edu/). A total of 735 participants from the ADNI cohort were identified as either Alzheimer’s disease (AD), mild cognitive impairment (MCI), or non-impaired (NI). Diagnostic criteria included Mini-Mental State Examination (MMSE) scores of 24-30 for NI and MCI patients and 20-26 for AD as well as a Clinical Dementia Rating (CDR) score of 0 for NI, 0.5 for MCI, and 0.5-1 for AD ^27^. In the current study, participants were allocated to groups (AD, MCI, or NI) based on their ADNI2 ‘current’ diagnoses. Participants’ age ranged from 71.3 to 76.5 across the groups and included a mix of males and females (Table 1). Clinical progression of cognitive impairment was based on participants’ most recent (by year) diagnosis in ADNI3. APOE genotype for ADNI participants was determined by blood sample ^27^.

**Table 1.** Characteristics of the included ADNI cohort participants.

|  |  | **AD+** | **AD-** | **MCI+** | **MCI-** | **NI+** | **NI-** |
| --- | --- | --- | --- | --- | --- | --- | --- |
| Total N |  | 209 | 100 | 121 | 162 | 33 | 110 |
| Sex (M:F:unknown) |  | 118:90:1 | 61:39 | 71:49 | 91:70:1 | 20:13 | 54:56 |
| Age (Average +/- SD) |  | 73.6 ± 0.5 | 76.5 ± 0.9 | 71.3 ± 0.7 | 73.3 ± 0.6 | 74.5 ± 1.1 | 74.6 ± 0.6 |
| APOE genotype | e2,e2 | - | 1 | - | 1 | - | 0 |
|  | e2,e3 | - | 7 | - | 19 | - | 22 |
|  | e3,e3 | - | 92 | - | 142 | - | 88 |
|  | e2,e4 | 4 | - | 4 | - | 1 | - |
|  | e3,e4 | 144 | - | 89 | - | 30 | - |
|  | e4,e4 | 61 | - | 28 | - | 2 | - |

“+” indicates APOE4 carriers and “-” indicates APOE4 non-carrier

### 2.2. CSF Proteomics

CSF proteome data accessed through the ADNI database was generated by the Neurogenomics and Informatics Centre at Washington University (https://neurogenomics.wustl.edu/) and the Cruchaga Lab at Washington University School of Medicine (https://cruchagalab.wustl.edu/). CSF proteome samples were analyzed using the SomaScan 7k assay. Data was normalized by SomaLogic including hybridization and median normalization and normalization to a reference using iterative Adaptive Normalization by Maximum Likelihood (ANML) ^28^. Following normalization, additional QC procedures were applied as described in Wang et al. ^28^. Protein levels are reported as Relative Fluorescence Unit (RFU). Although there was proteome data for 758 participants from the ADNI cohort in this dataset, APOE genotype was missing for 23 of these and were thus excluded from the current study (final n = 735). Protein hits in the CSF proteome file were mapped to Uniprot IDs using the package ‘SomaScan.db’ in R (v4.3.1). Proteins that were unable to be mapped to Uniprot IDs were removed prior to further analysis.

### 2.3. CSF AD Pathology Burden

To determine whether APOE4 carriers had a high level of AD pathology burden, we identified AD biomarker CSF data including Aβ_42_, total tau (t-tau), and phospho-tau181 (p- tau181). Data was similarly sourced through ADNI and generated by the Department of Pathology & Laboratory Medicine and Centre for Neurodegenerative Diseases Research at the University of Pennsylvania. As described elsewhere ^29^, Roche Elecsys immunoassays were used to detect Aβ_42_, t-tau, and p-tau181 in participant CSF samples according to the manufacturer’s instructions. Reference ranges for each analyte were (lower to upper limit) 200-1,700 pg/ml for Aβ_42_, 80-1,300 pg/ml for t-tau, and 8-120 pg/ml for p-tau181 ^29^.

### 2.4. Statistical Analyses

To perform CSF proteome profiling and identify proteins that may be driving between group differences, we employed our unique machine learning methods, as previously described ^30,31^. Here, we first use weight-of-evidence and information value for feature selection to identify a subset of proteins that are the strongest predictors (>0.3) of group differentiation. We then evaluated the predictive performance of this subset of proteins using classification and regression trees (CART). Importantly, the predictive performance of each protein was evaluated both independently and together with other proteins. This allowed us to identify proteins that may be themselves drivers of between group differences but also proteins that interact together to drive these differences. The CSF proteome dataset was split into 70% training and 30% held-out testing datasets. CART models were built, fine-tuned, and validated on the training dataset using five-fold cross-validation repeated five times. Models were fine-tuned using an automatic grid with 100 parameters. An important consideration in the current dataset was the presence of clear class imbalances across the groups (Table 1). More specifically, the n’s were unequally distributed between the group which can affect the validity and reliability of machine learning models like CART. To account for class imbalances, we used both oversampling and undersampling techniques to equalize the n’s across groups and report model performance metrics that are an average of both. All CART models were evaluated using the 30% held-out dataset. Performance was determined using several performance metrics including sensitivity, positive predictive value (PPV; also known as precision), specificity, negative predictive value (NPV), and area under the curve (AUC).

To identify whether APOE4 carriers were more likely to progress to MCI and AD over time, we obtained longitudinal clinical data for the 735 participants through the ADNI database (‘ADNI 4’ diagnosis). We used a Chi-squared analysis with *p* < 0.05 to compare the likelihood of progression between APOE4^+^ NI (NI APOE4 carriers) and APOE4^-^ NI (NI APOE4 non-carriers). We also examined whether the APOE4^+^ NI had a high level of AD pathology burden using a linear regression with age as a covariate and β and *p* values reported. All data analysis and figures were done in R v 4.3.1 (packages: ‘Information’, ‘caret’, ‘ggplot2’).

### 2.5. Functional Enrichment Analyses

We examined the relationship between identified proteins that were driving between group differences using protein-protein interaction (PPI) network analyses performed in NetworkAnalyst 3.0 ^32^. Here, generic PPIs were generated using IMEx interactome data ^33^. To examine the functional connectivity of the interactive proteins, we applied a Steiner Forest Network analysis that uses a fast heuristic Prize-collecting Steiner Forest (PCSF) algorithm. We then applied a first order network analysis to examine biological and molecular enrichment in Gene Ontology (GO) and Reactome pathways.

For brain region and cell type-specific enrichment analyses, the Human Protein Atlas was used ^34,35^ (https://www.proteinatlas.org/) (v23, Ensembl v109). Expression profiles for brain tissue were based on immunohistochemistry using tissue microarrays and data was included for all measured brain regions (cerebral cortex, caudate, hippocampus, and cerebellum) ^35^. Cell type specific data for brain and immune cells were based on single cell RNA-sequencing ^34^. For these enrichment analyses, normalized expression was used and heatmaps were generated by using min-max scaling and in GraphPad Prism (v.10.0.0 for Windows).

## 3. RESULTS

### 3.1. APOE4 carriers have a unique proteome signature irrespective of cognitive status

We first sought to identify differences in the CSF proteome of APOE4 carriers with AD (APOE4^+^ AD) relative to non-carriers with AD (APOE4^-^ AD). To do this, we performed an initial feature selection of the 6,082 proteins identified by the semi-targeted proteomics SomaScan 7k assay using weight-of-evidence and information value. This identified 1,534 proteins that had strong predictive power (>0.3) of between group differences (Supplementary Table 1). We then examined the ability of the 1,534 proteins, both independently and together, to predict APOE4^+^ AD relative to APOE4^-^ AD using CART. Two groups of proteins had a strong predictive performance (sensitivity >0.75). The first group was characterized by seven “stand-alone” proteins that each independently predicted APOE4^+^ AD and APOE4^-^ AD cases. Performance metrics for each of the seven independent proteins were all 1.0 for sensitivity, specificity, PPV, NPV, and AUC, respectively. The independent proteins included CCL25, CHCHD7, LRRN1, Otulin, S100A13, SPC25, and TBCA (Figure 1, Supplementary Table 2).

**Figure 1.**
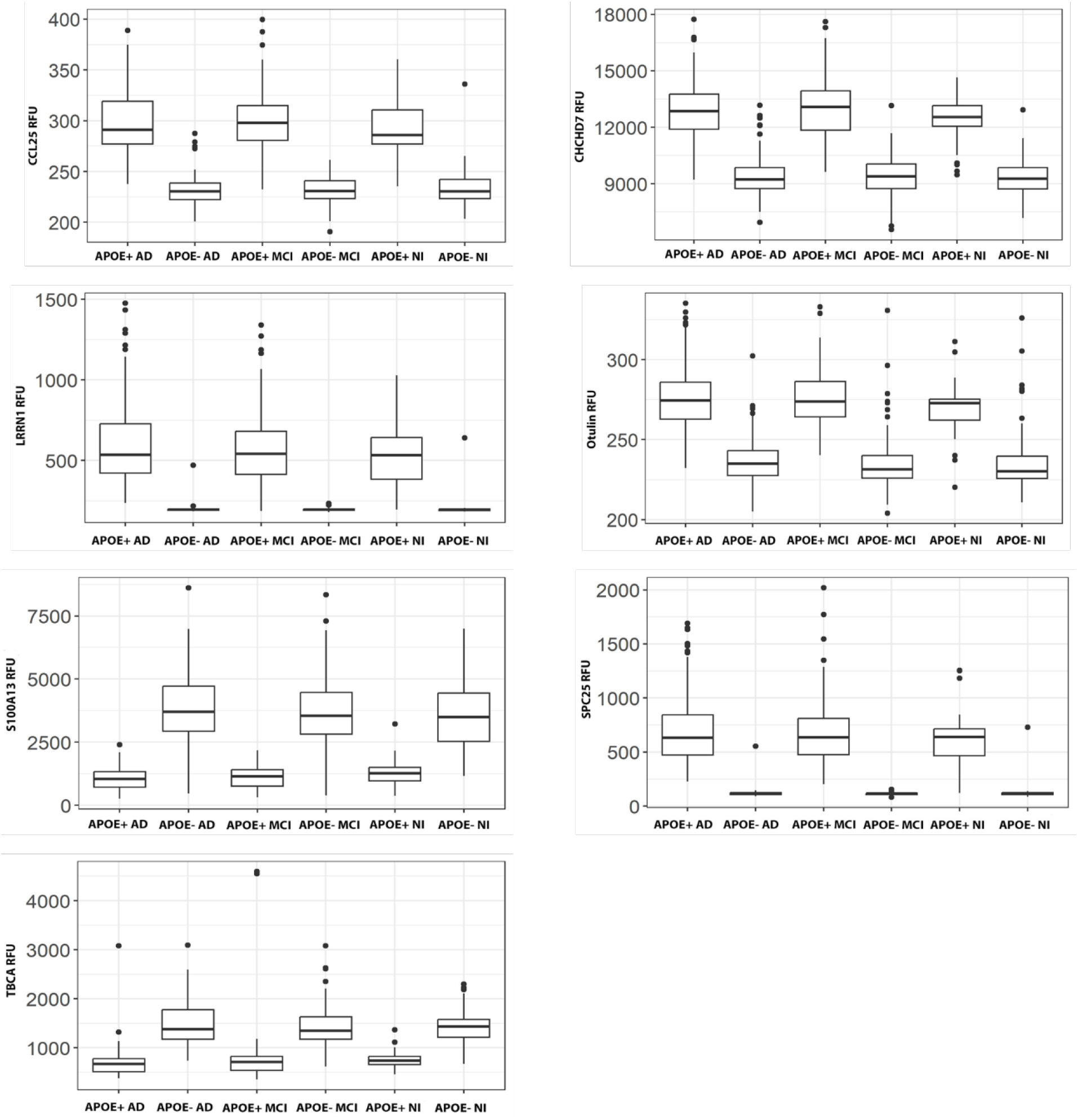
Box plots of the seven independent proteins that each predict APOE4 carriers irrespective of cognitive status. Plots show the relative fluorescent units (RFU) of each protein CCL25, CHCHD7, LRRN1, Otulin, S100A13, SPC25, and TBCA and highlight differences between APOE4 carriers and non-carriers. Abbreviations: AD: Alzheimer’s disease; MCI: mild cognitive impairment; NI: no impairment.

The second group of proteins was characterized by 50 proteins that interacted together to predict APOE4^+^ AD and APOE4^-^ AD (interactive proteins; Supplementary Table 3). These proteins similarly showed high CART performance metrics (sensitivity = 0.99, specificity = 0.74, PPV = 0.86, NPV = 0.98, AUC = 0.92), indicating that they were strong predictors.

We next examined whether this proteome signature of independent and interactive proteins was unique to APOE4^+^ AD or whether it was generalizable to other groups. To do this, we first tested whether our CART models and signatures could generalize to APOE4 carriers with MCI or NI (APOE4^+^ MCI or APOE4^+^ NI). CART models using either the independent or interactive proteins as predictors performed poorly and lost their predictive power (Table 2). We used CART to further test whether our proteins could differentiate between APOE4^+^ MCI and APOE4^+^ NI; similarly finding that our models were unable to do so (Table 2). This demonstrated that both the independent and interactive proteins are the same across all APOE4 carriers, independent of cognitive status (AD, MCI, or NI).

**Table 2.**
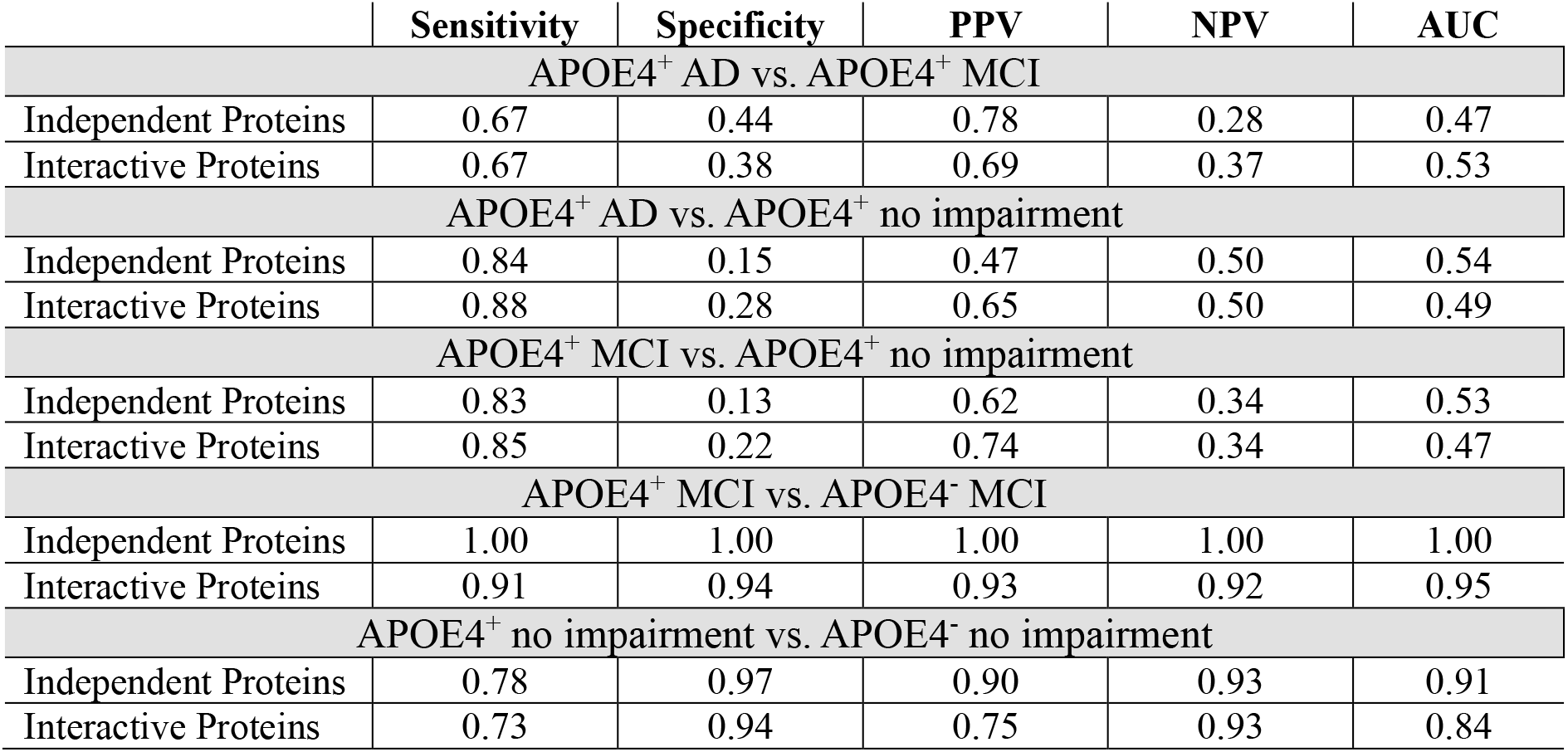
Performance metrics of CART models testing the uniqueness of the identified proteome signature to APOE4 carriers.

|  | Sensitivity | Specificity | PPV | NPV | AUC |
| --- | --- | --- | --- | --- | --- |
| APOE4<sup>+</sup> AD vs. APOE4<sup>+</sup> MCI |  |  |  |  |  |
| Independent Proteins | 0.67 | 0.44 | 0.78 | 0.28 | 0.47 |
| Interactive Proteins | 0.67 | 0.38 | 0.69 | 0.37 | 0.53 |
| APOE4<sup>+</sup> AD vs. APOE4<sup>+</sup> no impairment |  |  |  |  |  |
| Independent Proteins | 0.84 | 0.15 | 0.47 | 0.50 | 0.54 |
| Interactive Proteins | 0.88 | 0.28 | 0.65 | 0.50 | 0.49 |
| APOE4<sup>+</sup> MCI vs. APOE4<sup>+</sup> no impairment |  |  |  |  |  |
| Independent Proteins | 0.83 | 0.13 | 0.62 | 0.34 | 0.53 |
| Interactive Proteins | 0.85 | 0.22 | 0.74 | 0.34 | 0.47 |
| APOE4<sup>+</sup> MCI vs. APOE4<sup>-</sup> MCI |  |  |  |  |  |
| Independent Proteins | 1.00 | 1.00 | 1.00 | 1.00 | 1.00 |
| Interactive Proteins | 0.91 | 0.94 | 0.93 | 0.92 | 0.95 |
| APOE4<sup>+</sup> no impairment vs. APOE4<sup>-</sup> no impairment |  |  |  |  |  |
| Independent Proteins | 0.78 | 0.97 | 0.90 | 0.93 | 0.91 |
| Interactive Proteins | 0.73 | 0.94 | 0.75 | 0.93 | 0.84 |

We then tested whether the proteome signatures were able to differentiate between APOE4 carriers and non-carriers. CART models comparing APOE4^+^ MCI and APOE4^-^ MCI as well as APOE4^+^ NI to APOE4^+^ NI all had a high level of performance, similar to our initial models comparing APOE4^+^ AD to APOE4^-^ AD (Table 2). This indicated that the proteome signature is indeed specific to APOE4 carriers and, to further validate this, we performed a principal component analysis (PCA). This showed that there was no group separation when looking at all 6,082 identified CSF proteins (Figure 2a) but very clear group separation based on APOE4 status using our 57-protein (independent and interactive) proteome signature (Figure 2b). This effect was also visualized using a heat map based on measured protein levels (Figure 3), further highlighting that the CSF proteome signature was indeed unique to APOE4 carriers irrespective of cognitive status.

**Figure 2.**
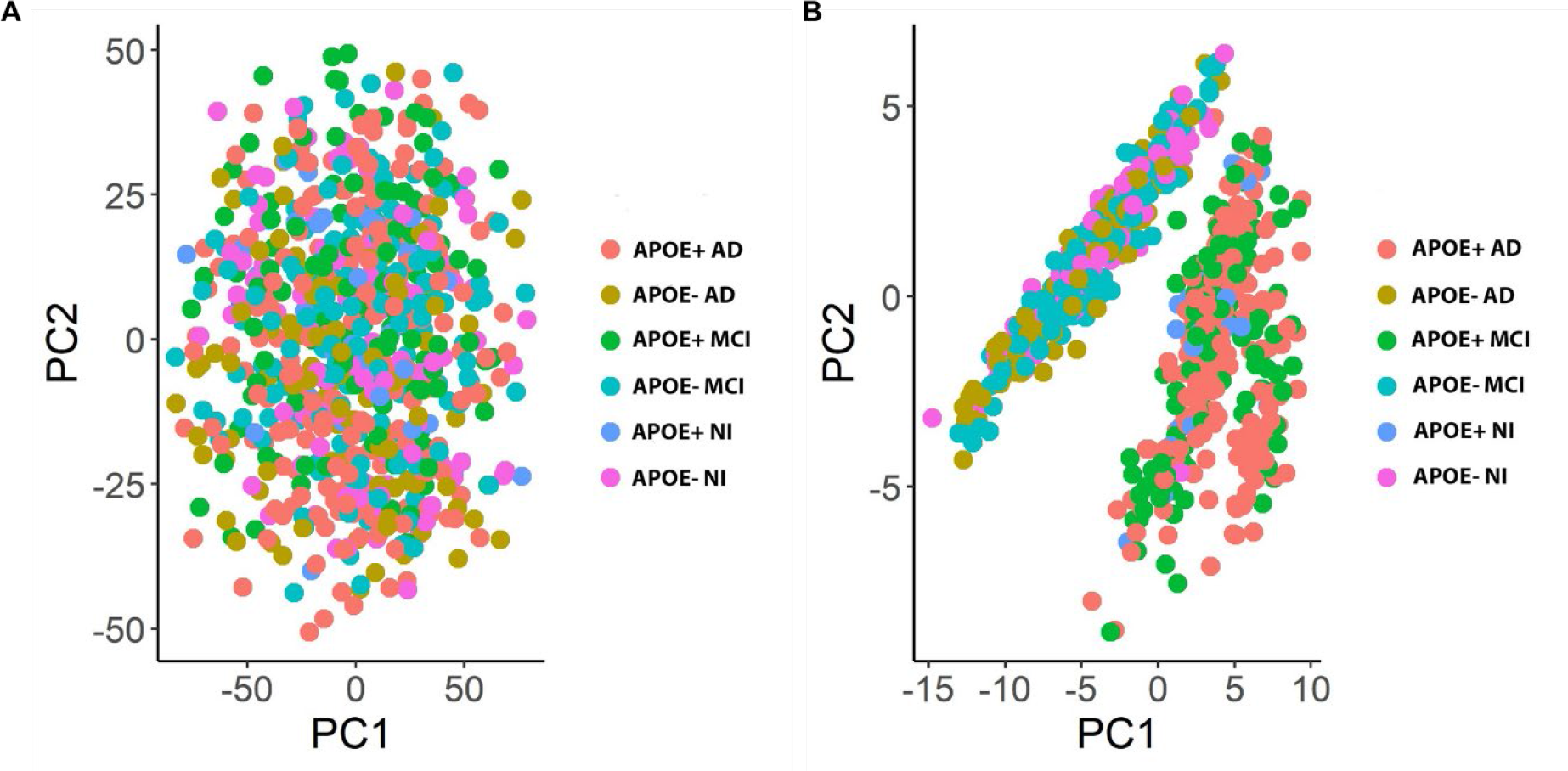
Principal component analysis (PCA) of CSF proteins across experimental groups. (**A**) PCA of all 6,082 proteins included in the present study shows no clear group separation based on cognitive status or APOE4 genotype. (**B**) PCA of the 57-protein proteome signature (including both independent and interactive proteins) identified as being unique to APOE4 carriers independent of cognitive status. Abbreviations: AD: Alzheimer’s disease; MCI: mild cognitive impairment; NI: no impairment.

**Figure 3.**
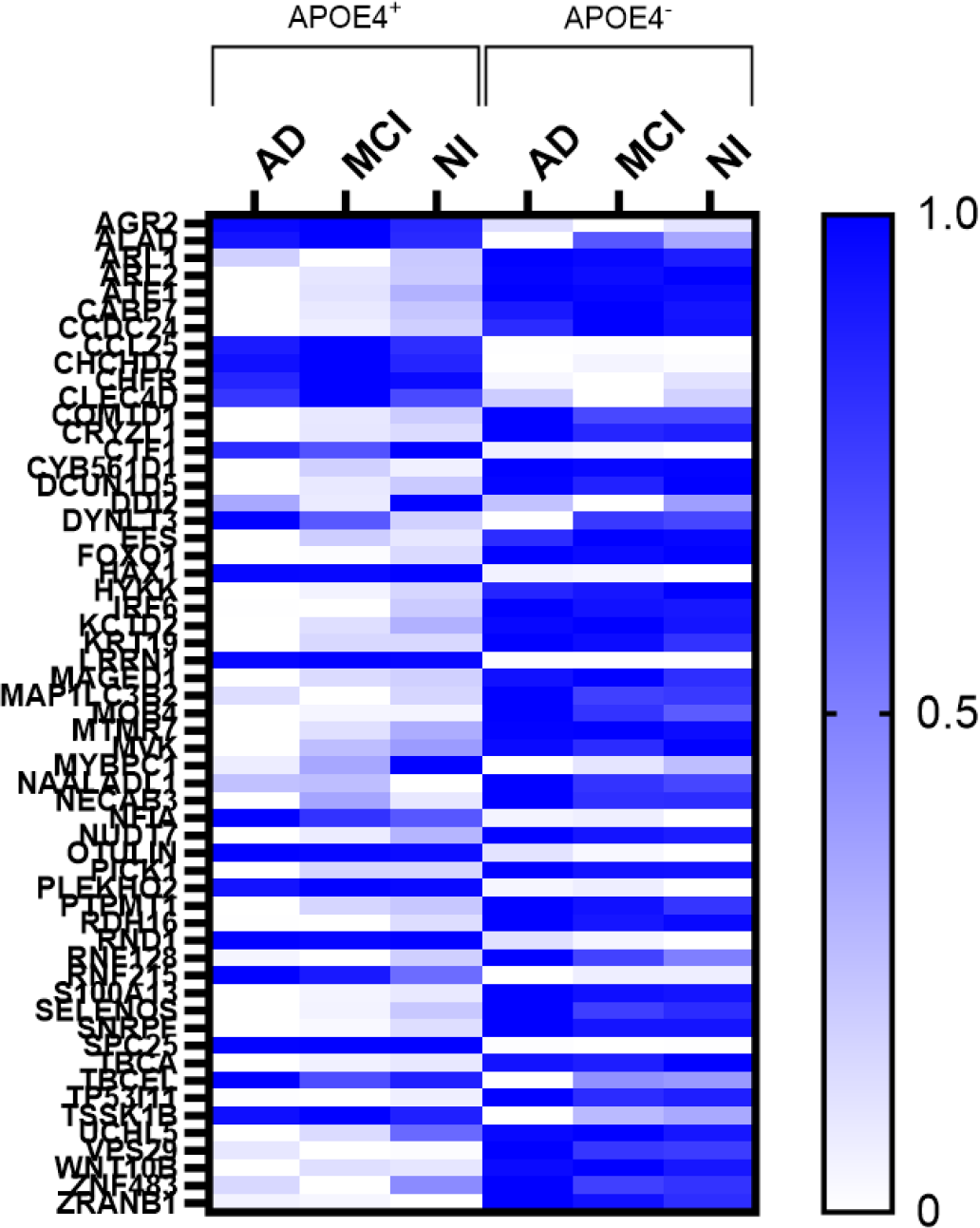
Heat map of the relative levels of 57 proteins (both independent and interactive) identified by CART as being unique to APOE4 carriers relative to non-carriers irrespective of cognitive status. Abbreviations: AD: Alzheimer’s disease; MCI: mild cognitive impairment; NI: no impairment.

### 3.2. The APOE4 proteome signature is independent of CSF AD pathology burden in non-impaired controls

Previous research has shown that APOE4 carriers may have a higher AD pathology burden relative to non-carriers, including, for example, higher cortical Aβ deposition ^36^ and increased tau spreading ^37^. An important consideration, therefore, was whether our APOE4- specific proteome signature was a consequence of increased AD pathology burden or occurs independently of this. To determine this, we compared CSF AD biomarker data for Aβ_42_, t- tau and p-tau181 for APOE4^+^ and APOE4^-^ NI participants using a linear regression. We also included age as a co-variate due to research showing that Aβ_42_ and tau levels also increase with normal aging ^38–41^.

Our analysis revealed non-significant main effects of APOE4 genotype on t-tau (β = −0.186, *p* = 0.795), p-tau181 (β = −0.166, *p* = 0.831), and Aβ_42_ (β = 0.158, *p* = 0.885). Additionally, there was no significant interactions between APOE4 genotype and age for t-tau (β = 0.005, *p* = 0.602), p-tau181 (β = 0.005, *p* = 0.594), or Aβ_42_ (β = −0.007, *p* = 0.625), indicating that the relationship between age and CSF AD biomarkers does not differ based on APOE genotype in NI participants. There was, however, significant main effects of age on t-tau (β = 0.013, *p* = 0.005) and p-tau181 (β = 0.015, *p* = 0.003), suggesting that t-tau and p- tau181 vary with age regardless of APOE genotype. Unlike tau, we did not find a significant main effect of age on Aβ_42_ levels (β = −0.006, p = 0.423).

### 3.3. APOE4 carriers are more likely to progress to cognitive impairment and AD

We next sought to identify whether the APOE4 proteome signature was pathogenic or benign by examining whether APOE4 carriers were more likely to progress to clinically diagnosed cognitive impairment over time. We found that APOE4^+^ NI or APOE4^+^ MCI participants in the current study were more likely to progress to either MCI or AD relative to participants without an APOE4 allele (14% vs. 5% progressed, respectively; X^2^ = 7.14, df = 1, *p* = 0.008).

### 3.4. Functional characterization of the APOE4 proteome signature

Finally, we sought to identify potential mechanisms underlying APOE4 carriers’ vulnerability to progression. Independent proteins (n=7) within the APOE4 CSF proteome signature were involved in T cell development, synapse assembly, innate immune responses, and cell division (Supplementary Table 3). We then performed a functional enrichment analysis on the interactive proteins (n=50) because they were interacting together to predict between APOE4 carriers and non-carriers. A Steiner Forest Network protein-protein interaction (PPI) analysis showed that 41/50 proteins formed a clear functional network (Figure 4).

**Figure 4.**
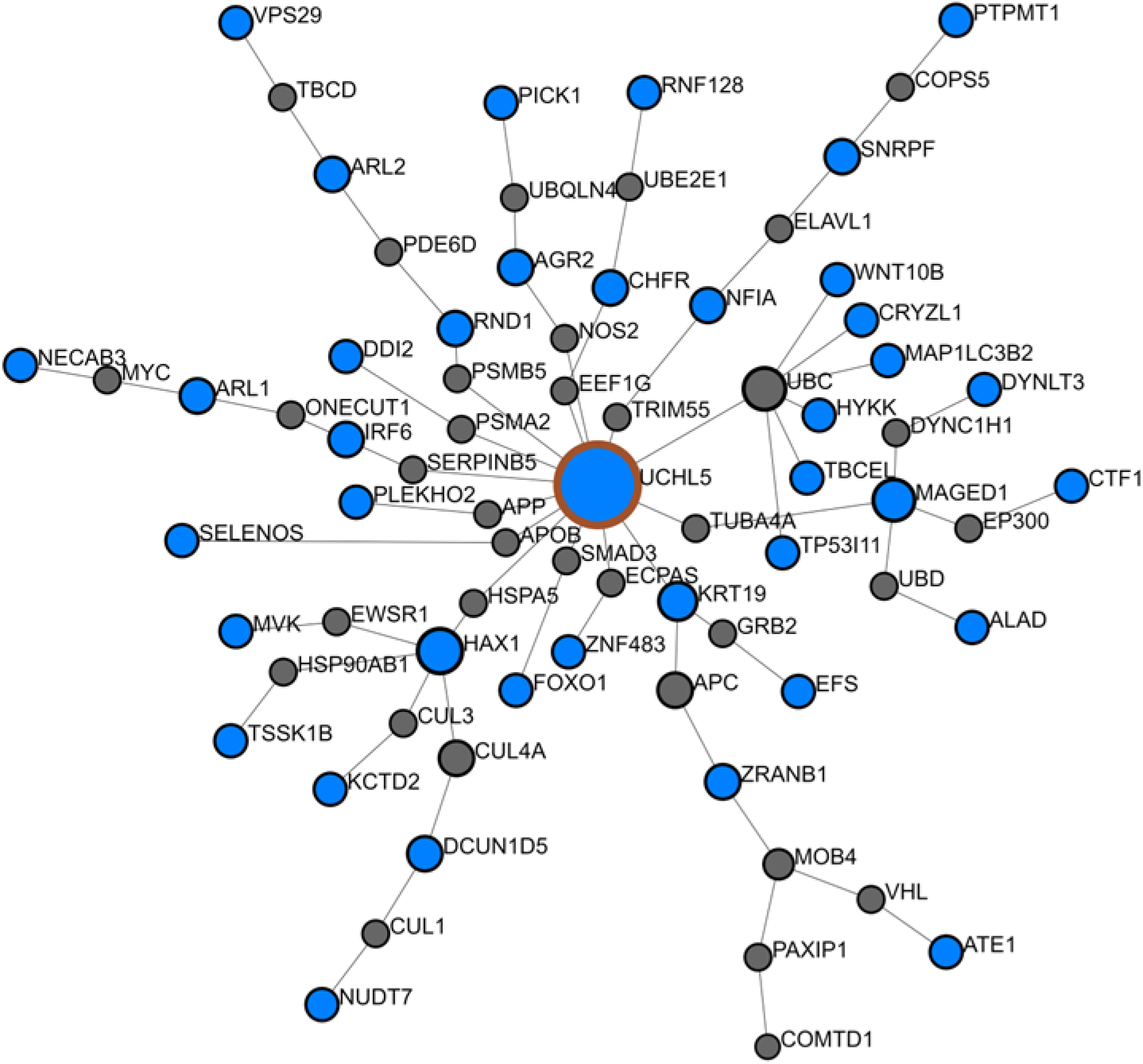
Steiner Forest Network protein-protein interaction (PPI) analysis showing functional connectivity between the interactive proteins (total n=41/50) in the APOE4 proteome signature. Blue nodes indicate proteins identified. Grey nodes indicate inferred proteins.

Further pathway enrichment analyses using GO indicated significant involvement in biological and molecular functions including catabolism, protein ubiquitination, mitosis, DNA damage response, ATP binding, and oxidative and cellular stress responses (all adj. *p* < 0.01; Supplementary Tables 4 and 5). Enriched Reactome signaling pathways included regulation of mitosis, immune system, apoptosis, inflammation and RNA and DNA regulation (all adj. *p* < 0.001; Supplementary Table 6).

To determine where our APOE4 proteome signature was enriched in the brain, we performed an enrichment analysis using immunohistochemistry microarray data from the Human Proteome Atlas ^35^. All 7 of the independent proteins were represented in the IHC microarrays however only 34/50 of the interactive proteins were represented. We found our proteome signature, including both independent and interactive proteins, was enriched in the caudate and cerebral cortex and, to a lesser degree, in the cerebellum (Figure 5). Interestingly, proteins within the signature were not especially enriched in the hippocampus (Figure 5).

**Figure 5.**
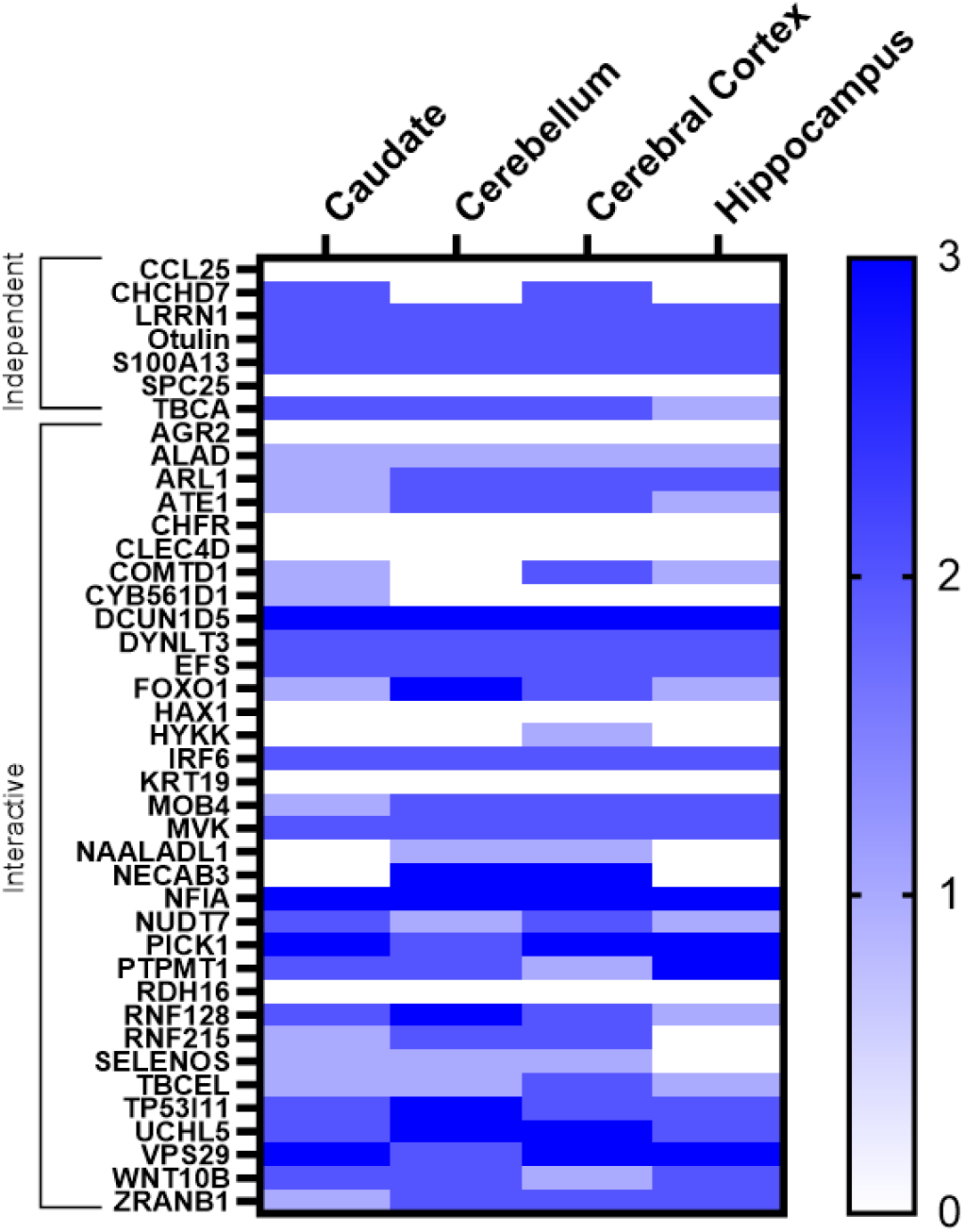
Enrichment across brain regions for the independent (n=7/7) and interactive (n=34/50) proteins within the APOE4 proteome signature based on confirmed immunohistochemistry microarrays from normal tissue in the Human Protein Atlas ^35^.

We then investigated cell type specific enrichment patterns for proteins within our APOE4 signature using single cell RNA-sequencing data from the Human Protein Atlas ^34^. In the brain, and limiting our analysis to only those proteins identified in as being expressed in the brain by IHC microarrays (Figure 5), we found the highest level of enrichment for both independent and interactive proteins in endothelial cells (Figure 6). Astrocytes and oligodendrocytes were also highly enriched for proteins within our APOE4 proteome signature, a finding in line with the immune and inflammatory pathways that these proteins are enriched for. Interestingly, this central immune dysregulation was also mirrored in the periphery. Single cell RNA-sequencing enrichment analysis for immune cell subtypes indicated that our APOE4 proteins (data found only for n=5/7 independent and n=36/50 interactive proteins) were especially enriched in macrophages and T cells and, to a slightly lesser extent, B cells (Figure 6).

**Figure 6.**
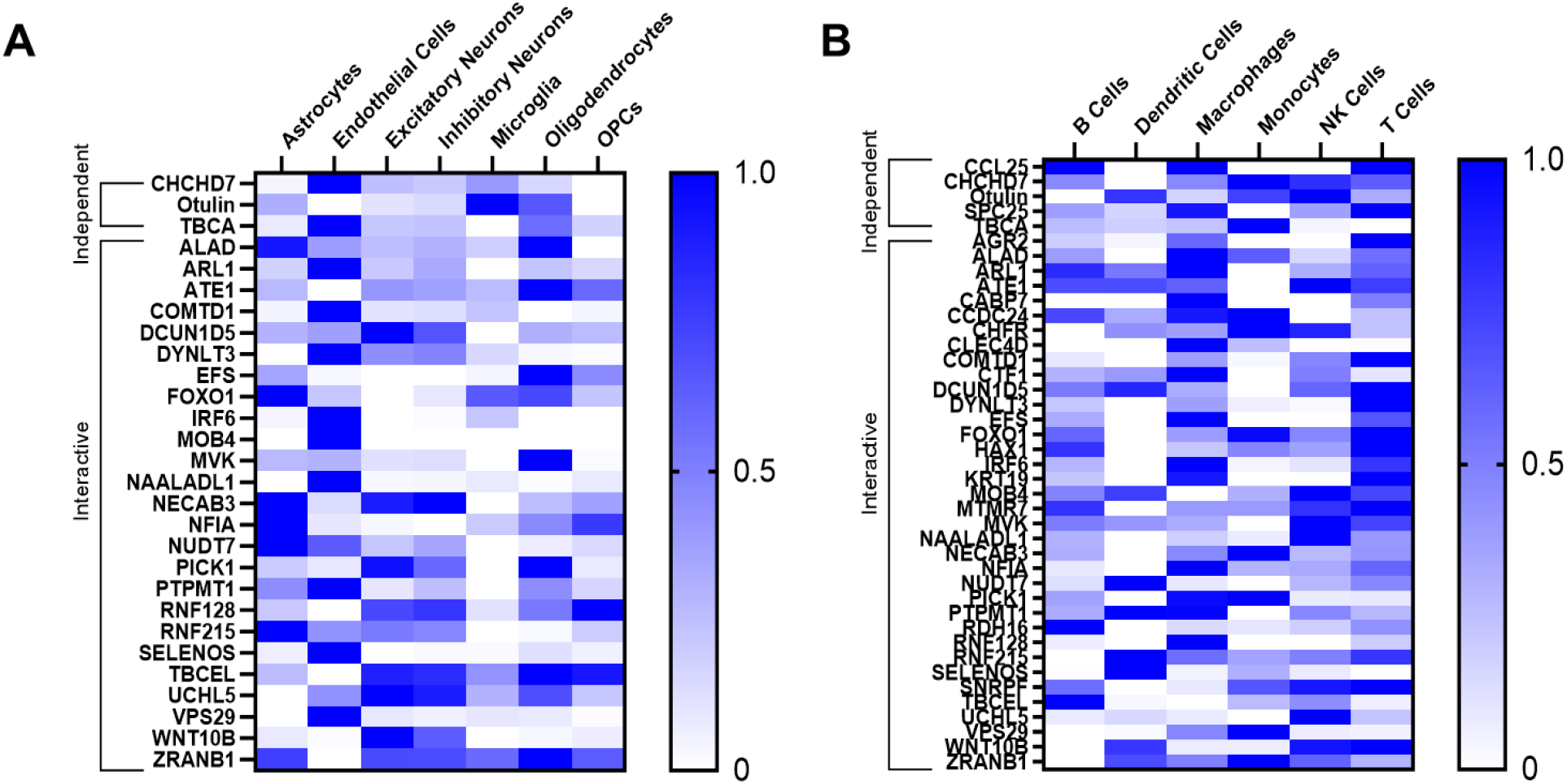
Cell type specific enrichment analyses for proteins within our APOE4 proteome signature across the brain and the periphery using single cell RNA-sequencing data from normal tissue from the Human Protein Atlas ^34^. (**A**) Enrichment for brain cell subtypes for the independent (n=3/7) and interactive (n=25/50) proteins shown to be expressed in brain tissue by IHC microarrays. (**B**) Enrichment across peripheral immune cell subtypes for independent (n=5/7) and interactive (n=36/50) proteins. Abbreviations: OPC: oligodendrocyte precursor cells.

## 4. DISCUSSION

While unique CSF and plasma proteins have previously been associated with APOE4^+^ AD patients ^23–25^, we expand on this existing literature by integrating machine learning and functional enrichment analyses to show that APOE4 carriers indeed have a unique CSF proteome signature consisting of seven independent and 50 interactive proteins. This signature was independent of cognitive status, as it was able to differentiate APOE4 carriers and non-carriers irrespective of whether they were not impaired, MCI, or AD. Importantly, we found that this protein signature was also independent of AD pathology burden. Using CSF AD biomarkers including Aβ_42_, t-tau, and p-tau181 we showed that in non-impaired healthy controls APOE genotype did not influence the levels of these AD biomarkers. This highlighted that the APOE4 CSF proteome signature in non-impaired controls was not due to the presence of increased AD pathology burden relative to cognitively normal APOE4 non-carriers. We also found that the APOE4 proteome signature was significantly associated with an increased risk of cognitive decline to MCI or AD over time. To our knowledge, ours is the first study to examine a large number of proteins (6,082) and demonstrate that there is a unique APOE4 CSF proteome that is not associated with AD pathology or cognitive status. Our findings are somewhat in line with previous research. A smaller study of less than 300 proteins used linear modelling to show that APOE4 genotype was associated with three unique CSF peptides when controlling for cognitive status ^26^. Further, only a few proteins were shown to be unique to APOE4 carriers when controlling for CSF t-tau, p-tau181, and Aβ_42_ ^26^.

Functional enrichment analyses of our identified 57-protein APOE4 proteome signature indicated significant enrichment for immune responses, inflammation, oxidative and cellular stress response, mitosis dysregulation, DNA damage, catabolism, protein ubiquitination, and synapses. Previous work has similarly highlighted a role for CSF markers of dysregulated immune processes and inflammation in APOE4 carriers. One study showed that APOE4 carriers with AD relative to non-carriers with AD have dysregulated complement pathway proteins that was independent of Aβ_42_ levels ^23^. Another, on the other hand, linked increased inflammatory cytokines, including IL-4, IL-6, and IL-8, to reduced Aβ deposition and preservation of cognitive function in APOE4 carriers with MCI and early AD ^42^. In a study of cognitively healthy APOE4 carriers, researchers found that APOE4 genotype was associated with decreased CSF TNFα levels ^43^. Importantly, in our study we found that the immune and inflammatory phenotype was reflected in both the brain and the periphery, suggestive of widespread, systemic changes in these processes. This is in line with previous research showing that APOE4 carriers have an increased innate immune response to challenges from lipopolysaccharide (LPS) and toll-like receptor stimulation ^44^. Further, we found that endothelial cells were significantly enriched for APOE4 proteins, suggestive of BBB dysfunction. A recent study showed that APOE4 carriers have clear BBB breakdown in the hippocampus and medial temporal lobe independent of Alzheimer’s disease pathology and prior to the onset of cognitive decline ^22^. This finding is supported by human iPSC-derived models of the BBB that show endothelial cells have a toxic gain of dysfunction, highlighted by proinflammatory states and cytokine release ^45^, dysregulated extracellular matrix and endothelial junction integrity ^46^, and cerebral amyloid angiopathy that’s driven by calcineurin-nuclear factor of activated T cells (NFAT) signaling ^21^. Future research should continue to use human-specific models to better understand the role that APOE4, and potential systemic immune dysregulation, plays in driving BBB breakdown and dysfunction.

An important finding of our paper is that the unique APOE4 CSF proteome signature was independent of both AD pathology and cognitive status. Future research would benefit from examining our identified proteins and enriched cell subtypes as potential precision medicine therapeutic targets for APOE4 carriers at risk of AD. Smaller studies have similarly shown that that APOE4 does not interact with tau/Aβ_42_ ratio in the CSF ^25^ and no association with CSF tau or p-tau181 ^47^. Previous studies, however, have found that APOE4 carriers tend to have reduced CSF Aβ_42_, even when they are cognitively normal ^47^. One reason for this disparate finding may be that the previous study did not explicitly control for age although do show that CSF Aβ_42_ continue to reduce with increasing age in APOE4 carriers. This highlights the importance of controlling for age when looking at markers of neurodegeneration to exclude age-related increases that may not necessarily be pathological.

Our study has limitations. First, the lower n and use of machine learning for proteome profiling precluded examining potential differences between homozygous and heterozygous APOE4 carriers. As outlined in Table 1, there were 61 homozygous carriers in the APOE4^+^ AD group whereas there were 28 and 2 in the APOE4^+^ MCI and APOE4^+^ NI groups, respectively. As documented extensively in the literature, there is a clear relationship between the number of APOE4 alleles and AD risk; with heterozygosity associated with a two to four times increased risk and homozygosity associated with a risk of 15 times or greater risk of AD, although this can vary depending on sex and ethnicity ^48,49^. Further, there is a growing argument that APOE4 homozygosity may represent a different form of AD that is more closely aligned to autosomal dominant AD ^50^. Although there may be differences in our APOE4 proteome signature between heterozygous and homozygous carriers, this is unlikely. Our CART models performed strongly and similarly across all performance metrics irrespective of the relative distribution of APOE4 genotypes between the groups. This shows that the presence or absence of homozygous APOE4 patients did not affect our models’ ability to use the proteome signature to predict APOE4 carriers, suggesting that all APOE4 carriers have this unique signature. Future research, however, should continue to source larger numbers of homozygous APOE4 carriers to further confirm this finding. A second limitation of our study was an inability to examine males relative to females, again due to lower n’s and our use of machine learning. The relative distribution of males to females did vary between the groups, ranging from as high as 61% male in the APOE4^-^ AD group to as low as 49% in the APOE4^-^ NI group (Table 1). Sex differences in the effects of APOE4 have been shown previously. For example, female APOE4 carriers with MCI have faster memory decline relative to APOE4 males with MCI ^51^, are more likely to progress to further cognitive decline and AD ^52^, and an interaction between menopause and APOE4 in females that contributes to greater lifetime AD risk ^53^. Again, however, it’s important to note that our CART models showed similar performance metrics across all comparisons irrespective of the relative distribution of males and females within the group. This strongly suggests that our unique APOE4 proteome signature is independent of sex although further confirmatory work is needed.

In conclusion, we found a CSF proteome signature that was unique to APOE4 carriers. This signature was independent of cognitive status, AD pathology as measured by CSF AD biomarkers, and was associated with an increased risk of future cognitive impairment. This suggests that this proteome signature may underly the increased risk of cognitive decline and AD in APOE4 carriers. In addition to being implicated in the BBB, our APOE4 proteins showed enrichment for the immune system and inflammation. From a precision medicine perspective, this suggests that immunomodulation and treatments targeted at improving the BBB may offer the best treatment strategies for APOE4 carriers at risk of AD.

## ACKNOWLEDGEMENTS

The authors are grateful to the Alzheimer’s Disease Neuroimaging Initiative for providing the data. The authors are thank the Neurogenomics and Informatics Centre at Washington University (https://neurogenomics.wustl.edu/) and the Cruchaga Lab at Washington University School of Medicine for generating the CSF SomaScan 7k assay proteome data (https://cruchagalab.wustl.edu/) for the ADNI cohort. We also thank the Department of Pathology & Laboratory Medicine and Centre for Neurodegenerative Diseases Research at the University of Pennsylvania for providing the Roche Elecsys AD CSF biomarker data for the ADNI cohort. Finally, the authors thank all the patients and their families for their involvement in ADNI.

## CONSENT STATEMENT

The ADNI cohort study was approved by the institutional review boards of the participating ADNI centers, and all participants provided informed consent.

